# An integrated multiscale, multicellular skin model

**DOI:** 10.1101/830711

**Authors:** Ryan Tasseff, Boris Aguilar, Simon Kahan, Seunghwa Kang, Charles C. Bascom, Robert J. Isfort

## Abstract

Skin is our primary barrier to the outside world, protecting us from physical, biological and chemical threats. Developing innovative products that preserve and improve skin barrier function requires a thorough understanding of the mechanisms underlying barrier response to topical applications. In many fields, computer simulations already facilitate understanding, thus accelerating innovation. Simulations of software models allow scientists to test hypothesized mechanisms by comparing predicted results to physical observations. They also enable virtual product optimization, without physical experiments, once mechanisms have been validated. The physical accessibility and abundant knowledge of skin structure makes it a prime candidate for computational modeling. In this article, we describe a computational multiscale multicellular skin model used to simulate growth and response of the epidermal barrier. The model integrates several modeling styles and mathematical frameworks including ordinary differential equations, partial differential equations, discrete agent-based modeling and discrete element methods. Specifically, to capture cell biology and physical transport, we combined four distinct sub-models from existing literature. We also implemented methods for elastic biomechanics. Our software implementation of the model is compatible with the high-performance computing simulation platform Biocellion. The integrated model recapitulates barrier formation, homeostasis and response to environmental, chemical and mechanical perturbation. This work exemplifies methodology for integrating models of vastly different styles. The methodology enables us to effectively build on existing knowledge and produce “whole-system” tissue models capable of displaying emergent properties. It also illustrates the inherent technical difficulties associated with the mounting complexity of describing biological systems at high fidelity. Among the challenges are validation of the science, the mathematical representations approximating the science and the software implementing these representations. Responsibility for a discrepancy observed between *in silico* and *in vitro* results may as easily lie at one of these three levels as at another, demanding that any sustainable modeling endeavor engage expertise from biology, mathematics and computing.

## 1 Introduction

### 1.1 Model Summary

We have developed a computational multiscale MultiCellular Skin Model (MCSM) for simulation of epidermal skin barrier formation, homeostasis and response to environmental, chemical and mechanical perturbation (Fig. 1). Our modeling strategy was to integrate four distinct models, mostly specific to skin, that had previously been validated and described in the literature. The first of these, a three-dimensional, agent-based model of skin with cells represented by discrete interacting spherical agents [1], provided the foundation. However, spherical agents cannot represent the flattening of cells as they rise toward the stratum corneum nor their elasticity when subjected to pressure. For these purposes, we replaced the spherical representation with a novel elastic ellipsoid representation that forms elastic adherence junctions. Secondly, for transport and diffusion of molecular species in the extracellular space, we employed a skin penetration model based on continuous Partial Differential Equations (PDEs) [2]. A continuum PDE representation was also employed as a third model to capture hydration effects and water transport, which modulates cell swelling and Trans-Epidermal Water Loss (TEWL) -a common clinical measure of barrier function [3]. The fourth model represents intracellular biomolecular processes in individual cells using Ordinary Differential Equations (ODEs). Specifically, regulation of cell proliferation was represented by a simplified model of the eukaryotic cell-cycle [4] that runs independently and in parallel across multiple cellular agents. The integration of these four mathematical models was achieved using software that runs on the high-performance computing platform Biocellion. Biocellion was designed to simulate macroscopic-scale living-system models at cell-resolution [5]. This integrated “whole-system” MCSM was able to recapitulate dynamic and emergent system behaviors that its constituent models cannot.

**Figure 1:**
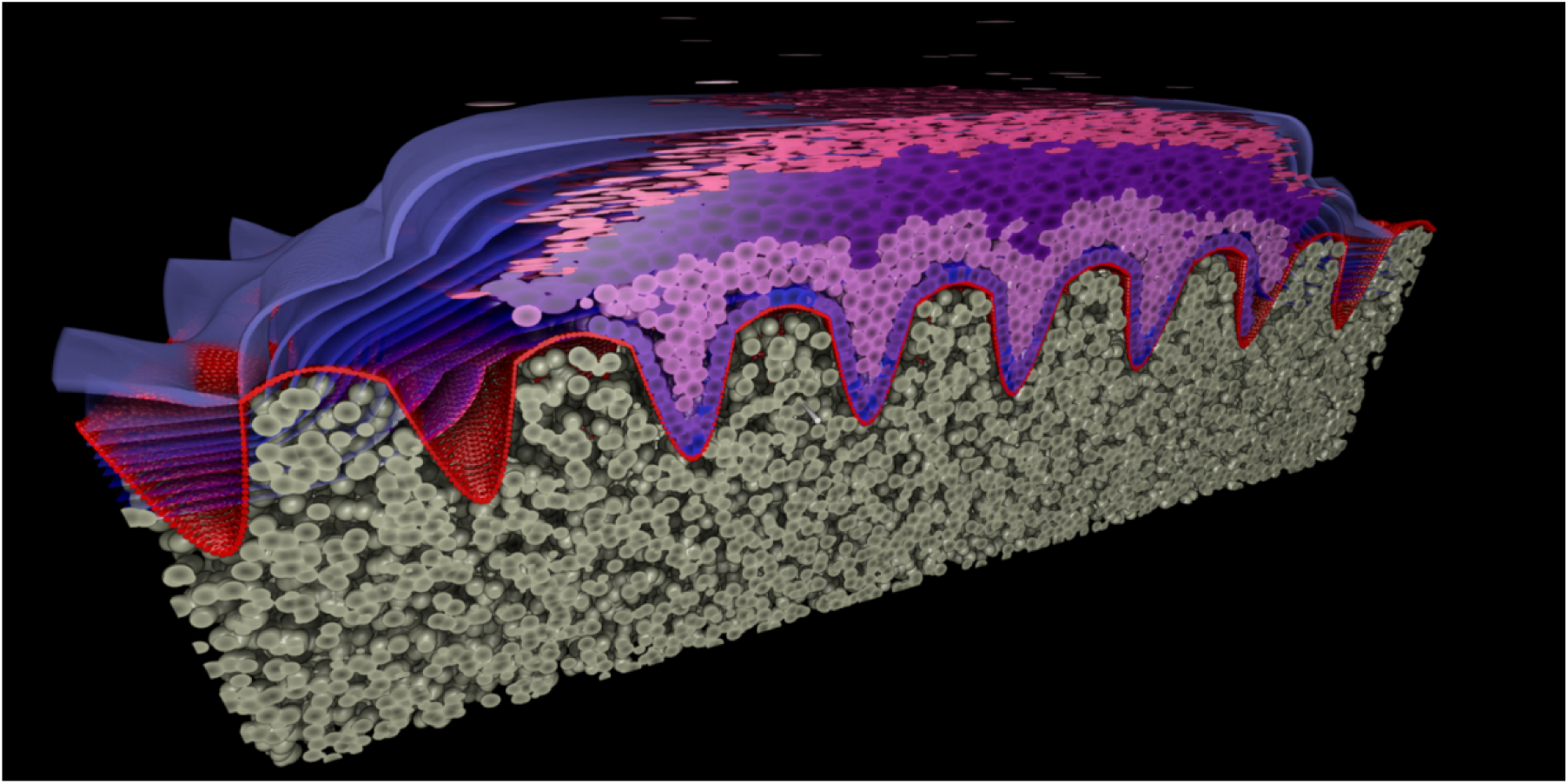
Example cross-section from a simulation of barrier homeostasis and water transport represented as blue isotherms. The image is cosmetically enhanced.

### 1.2 Motivation

There are several motivating factors for this work. The methodology itself was of interest for (i) the integration of knowledge and specific models across different disciplines and frameworks; (ii) the study of phenomena that arise from this integration including hard to capture feedback between the models and even more novel emergent properties; and (iii) the communication of results not only as quantitative metrics but also as visualizations of three dimensional tissues.

There was good reason for the consideration of skin, and particularly the epidermis. Skin is the largest human organ, but it also one of the simplest, and its primary physiological purpose as a regenerating barrier is manifested at scales less than a millimeter, where as organs like the heart, lungs and brain function at much larger length scales. It is also a more accessible system for clinical study compared to other tissues, and organotypic skin models can be grown in the laboratory directly from the underlying cell types [6]. These features simplify the modeling effort.

From the perspective of the Consumer Packaged Goods (CPG) industry, skin and epithelial tissues interact with nearly all products. Thus, a skin model would be very useful in designing new products, improving the function of existing products and helping the customers understand the benefits of products.

### 1.3 Related Work

The importance of skin accounts for the many prior and ongoing efforts to model skin barrier formation and homeostasis. Underlying these efforts are diverse modeling methodologies. These methodologies can be broadly classified in continuous and discrete models.

In continuous models, biological knowledge on the cellular level needs to be formulated in terms of continuous equations describing the impact on the tissue. Examples include Compartment and Finite-Element (FEM) models of skin. Compartment models are spatially coarse grained and represent physiological components of skin by a set of simplified, homogeneous compartments. Chemical and biological species can move between compartments. Their interaction is typically modeled using differential equations. Although physicochemical models that use PDEs to capture mass transport could be considered compartment models, there are also those that represent physiological processes. These typically assume well-mixed compartments and employ ODEs and standard reaction kinetic formulations (e.g. mass-action, Michaelis-Menten, empirical Hill functions).

Such physiological models can be very specific as exemplified by pharmacodynamic and pharmacokinetic models that capture an established enzymatic reaction involving a drug and its movement and metabolism using measured kinetic parameters. Complex biological processes can also be modeled such as predicting the toxicity of a material or assessing risk of allergic reaction [7]. Increasingly complex processes tend to require empirical functions with abstract parameters that do not have a physical representation. These are fit using data and can cause statistical issues with increasing dimensionality. ODE models utilize a well-established toolbox from extensive application in nonlinear dynamics and modeling engineering processes. These models do not capture biomechanics. FEM has been used to predict, in contrast, the biophysical properties of skin, such as elasticity and wrinkling, by representing the skin barrier as a two [8] or three dimensional biomass continuum. Use of FEM is attractive due to its proven use over 70 years in aeronautic, automotive and other engineering industries and its consequently mature software base. These models can include biological processes such as tissue growth, but only as it pertains to the mechanics of the system. A detailed review is provided by Limbert [9].

In contrast, discrete agent modelling allows direct implementation of such knowledge and the manifest tissue behavior naturally emerges. Overcoming this limitation, ABM has proven capable of predicting tissue-scale properties from those of individual cells by representing each skin cell as an autonomous three-dimensional agent whose behaviors are dictated by explicit mechanistic rules [1, 10]. The attraction of ABM is the correspondences it makes both of three-dimensional simulated outcomes to those of physical experiment and of the explicit rules to knowledge or hypotheses. Virtual experiments simulating ABMs mirror physical experiments, so the cause of divergent results can be mechanistically traced backwards to incomplete or incorrect assumptions around an individual cell’s behavior. Unfortunately, while conceptually simple, every instance of ABM is as complex as its rule set, requiring custom software development whenever a modeler introduces a new rule.

Some ABM software systems aim to mitigate this challenge by providing implementations of rules relevant to a specific biological domain of application. EPISIM [11] is a software package promising ease of use in modeling epithelial tissues. While applications can provide easy development for some processes, it can still require significant development efforts for others. For example, to recapitulate skin barrier growth from stem cells to homeostasis and TEWL over varying humidity, the authors needed to extend EPISIM with addition of a biomechanical ellipsoidal cell model [12]. Extension and refinement of models by software developers to incorporate new kinds of knowledge as their predictive capability and accuracy increase is unavoidable.

### 1.4 Approach

To facilitate model extension and refinement, we utilized the Biocellion modeling and simulation platform [5]. Similarly to EPISIM, Biocellion supports representation of each cell as an agent. A key difference is that while Biocellion is tuned to simulate biological systems versus more general ABM, the platform is not specialized to any particular biological domain. On the one hand, this means more software development must be done upfront by a modeler to use the platform. On the other, the platform imposes fewer constraints on modeling, easing extension and refinement. At this time, we believe there is not a sufficient example of model utility that would allow us to expressly rule out any particular physical/biological concept, mathematical framework or numerical implementation at the outset. Nor do we have a good understanding of the trade-off between model/process simplification and utility. Therefore, we opted for flexibility over easy-of-use for the development of a base model that may have broad future application.

The current work exemplified the benefits of this freedom. Here, we incorporated detailed mechanics that went beyond the flattening of keratinocytes and captured rotations and elastic deformation. Accurately modeling the elasticity or wrinkling of tissue as a function of cell properties, for example, requires the cells to rotate and deform. The Biocellion platform identifies each agent as merely a point (or some finite set of points) in three dimensions, and makes no assumptions about shape and interactions between adjacent agents. For this reason, we were able to integrate cell mechanics including rotation and deformation into the model (in fact, into reusable modeling software libraries) without changing the Biocellion platform itself.

Our overall approach is to incrementally construct a “whole-system” MCSM made actionable through computer simulation to predict response to biological, environmental, chemical and mechanical interventions as a function of the rules the model expresses. To provide needed context, we will elaborate briefly on the Biocellion platform in Section 2 before transitioning to how we have modeled the mechanics of the skin barrier in Section 3. In Section 4, we elaborate on the underlying biological sub-models and how they are integrated into the Biocellion whole-system model. Section 5 presents a number of simulation results for mechanical sub-models, for growth to homeostasis using the whole-system model, and for a few interventions. Finally, we discuss these results and future directions in Section 6.

## 2 Simulation Platform and Framework

### 2.1 Biocellion

Human tissues are composed of many cells, each of which is a complex organism. Cells interact with other cells and their micro-environment by migrating, adhering to or pressuring neighboring cells or abiotic structures; proliferating, differentiating or dying; and producing or consuming extracellular molecules. Complex tissue, such as skin develop as cells differentiate to perform specialized functions. The bulk or phenotypic properties we observe at the macroscopic scale emerge from these underlying interactions and behaviors. It is therefore critical to integrate our knowledge of individual cell behaviors and the interactions among cells and their microenvironment for the study of complex tissues and phenotypes.

Modeling these systems may become computationally prohibitive, particularly when systems are scaled to clinically relevant time and length scales that can manifest phenotypes of interest. Biocellion, is a simulation platform for multicellular biological systems that provides the efficient numerical solvers and high-performance computing solutions necessary to reach appropriate complexity and scale. The current version of Biocellion is 1.2 and is available at Biocellion.com. For this work, we employed several advancements in the mechanical description of cells and their physical interactions. This allowed us to also simulate the important biomechanical features of skin.

### 2.2 Framework Overview

Briefly, Biocellion maps a cell to a discrete agent. Discrete agents can also represent non-cellular features such as basement membrane. Each discrete agent has associated variables representing its position and state. They can be positioned anywhere within the simulation domain and are not restricted to a predefined grid. These variables are updated based on user provided rules. Updates of cellular state and mechanical interactions between cells are made with the largest, or ‘baseline’, time step of 10s. However, baseline time steps could be further broken down when computing more advanced mechanics as described below. Ordinary Differential Equations (ODEs) are used to solve dynamic systems within individual cells and update state variables. Here we implemented the Intel ODE solver and a time step of 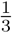 baseline.

Biocellion imposes a 3D cubic grid to the environment. The grid spacing was 50*µm*. Partial differential equations (PDEs) were used to model environmental state changes within grids due to molecular diffusion between grids. We implemented the CHOMBO PDE solver package, and we used a ‘grid’ time step of 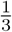 baseline. This is the time scale at which PDE and other grid box parameters can be changed. It was possible to have a smaller time step specifically for the PDE solver, e.g. to improve computational stability, but the grid and PDE time steps were the same. Grids were spatially divided into sub-girds and adaptive mesh refinement was used to provide increased or decreased resolution for the concentration gradients when appropriate. By default PDE parameters were constant for all sub-grids in a grid box unless otherwise stated.

## 3 Mechanical Interactions

Biocellion 1.2 was based on spherical agents. For this work we use quaternion representation of ellipsoids. We also account for deformation and improved cell-cell adhesion junctions. In the MCSM produced here, rotation is less critical, but may still play a role in specific mechanical perturbations. More importantly, the ellipsoidal representation better captures tissue morphology. The formation of flexible yet breakable junctions between cells are a critical aspect of cellular organization and mechanical properties of tissues. Finally, deformation plays a role in cellular morphology. Taken together, these upgrades allow us to better capture the interplay between mechanics and biology, and they bring us a step closer to simulating emergent behaviors; for example, the emergence of tissue level biomechanics.

### 3.1 Mechanical Description of Cells

Each cell was represented using two ellipsoids, one for the undeformed configuration (free of mechanical stress) and another for the deformed configuration (under mechanical stress). The later represented the actual cell shape. Both ellipsoids had the same orientation but their principal semi-axis lengths could differ.

Some cells could be spheres in their undeformed configuration; yet, their deformed shapes deviated from spheres immediately following cell division: a cell that divides vertically becomes shorter in the body-fixed z direction. A cell that divides horizontally shortens its principal semi-axis lengths in the body-fixed x and y directions. Each of these new bodies were assigned a spherical undeformed configuration after dividing, which would allow them to recover.

Typically cell growth increased the body-fixed horizontal principal semi-axis lengths without changing the ellipsoid height in the vertical direction in the undeformed configuration while size increases due to hydration acted in the vertical direction. Note that a cell’s body-fixed vertical direction did not always coincide with the body-fixed z direction if cells rotated significantly.

### 3.2 Equation of motion

#### 3.2.1 Newtonian Dynamics

Assuming that the mass (*m*) and body-fixed inertia (*I*) of agents remained constant during the simulation, the equations of translation and rotation for the *i*th agent were:

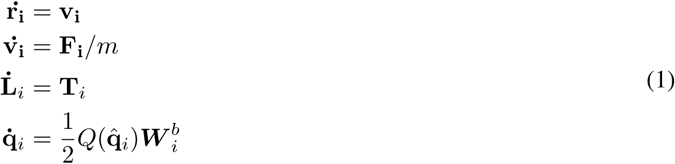

Where **r**_*i*_, **v**_*i*_, and **F**_*i*_ were the position, velocity, and total applied force on the center of agent *i*, respectively. Similarly, **L**_*i*_ and **T**_*i*_ were the angular momentum and the total torque on agent *i* in the reference frame. 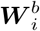 was the quaternion form of the angular velocity in the body-fixed frame, 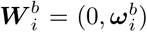, and 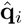 was unitary quaternion that define the rotation of the body-fixed rotation with respect to the reference frame, and:

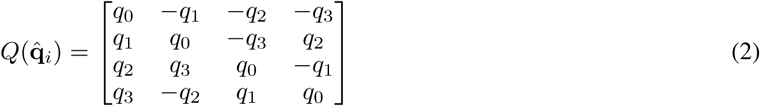

Using quaternions it was easy to rotate a vector from the body-fixed to the reference frame, for example to transform the momentum to the body-fixed frame we simply used 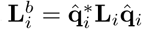. Moreover, 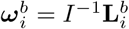.

### 3.3 Shoving and Adhesion

#### 3.3.1 Cell-Cell Contact

To compute shoving forces between a pair of ellipsoids we first computed the contact point (see Figure 2) using the geometric potential method [13], in which the contact point was defined as the midpoint between two points on ellipsoids in ellipsoids 1 and 0, *Point On Ellipsoid 0* and *Point On Ellipsoid 1* in Figure 2) [14]. To compute *Point On Ellipsoid 0* we minimized the following functional:

**Figure 2:**
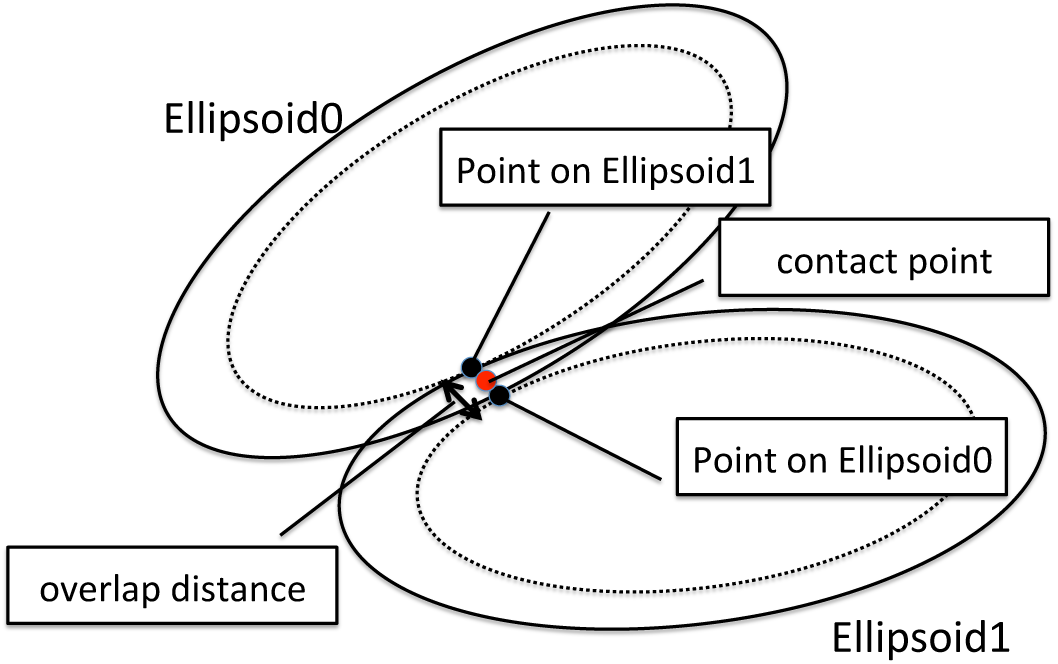
Consider two ellipsoids. If we scale *Ellipsoid 0* fixing the center, we can find the point the scaled *Ellipsoid 0* first barely touches *Ellipsoid 1*. This point is *Point on Ellipsoid 1*. Similarly, we can find *Point On Ellipsoid 0*. The contact point is defined as the midpoint of the segment joining the two Points.

**Figure 3:**
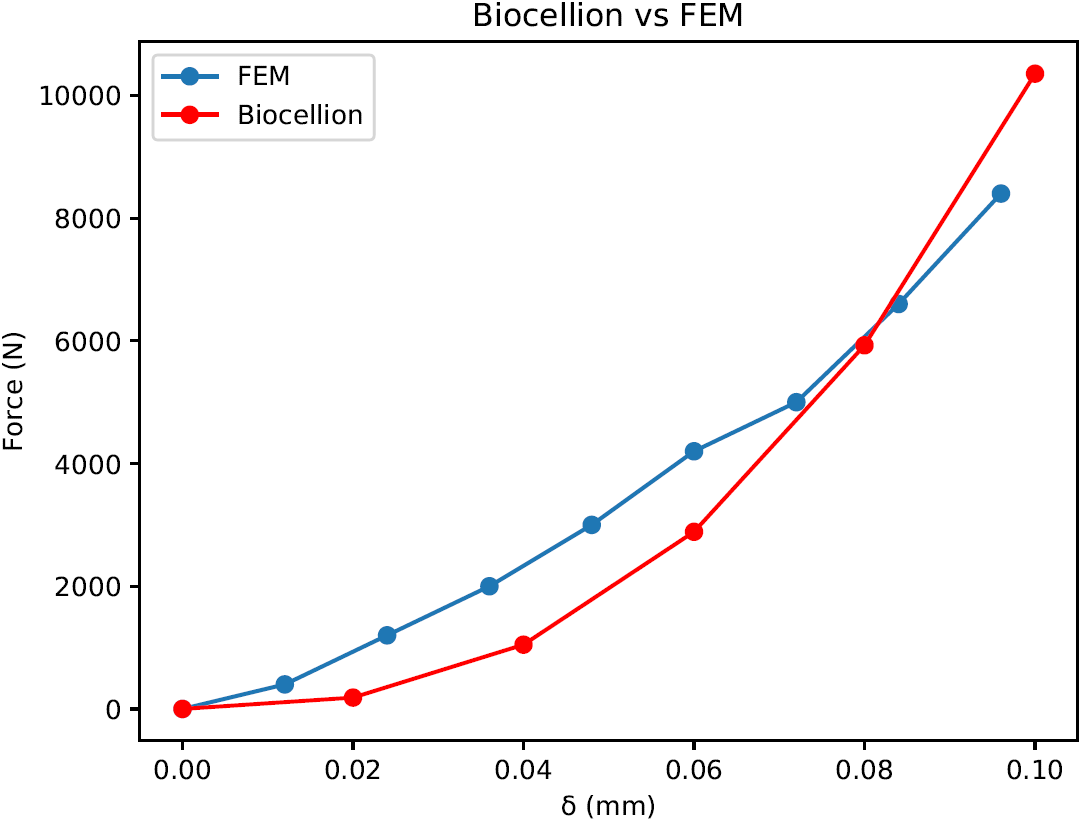
The normal contact force between two spheres computed with Finite Element method (FEM) and with Hertz theory implemented in Biocellion. The radii of the two cells are 5mm and 8mm, and the Young modulus and Poisson ratio are 2.110^1^2Pa and 0.3 respectively. The FEM based forces were obtained from Figure 4 of [17].

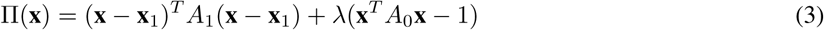

Where *A*_0_ and *A*_1_ were the 3D matrices that define ellipsoids 0 and 1 respectively, **x**_1_ was the center of ellipse 1. We assumed all the positions were defined with respect to the center of ellipsoid 0. *λ* was the Lagrange multiplier that ensured *x* belonged to the surface of ellipsoid 0. By making ∇Π(**x**) = 0 we obtained the following equation that defined *Point On Ellipsoid 1*:

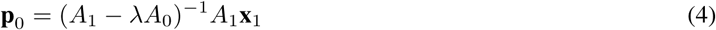

We solved the equation 4 iteratively by doubling the value of *λ* until the 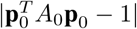 became smaller than a desired precision. The potential *Point On Ellipsoid 1*, **p**_1_ was computed with same procedure.

Then we computed the overlap distance *δ*_*n*_. The most straightforward approach to computing the overlap distance was to measure the distance between *Point On Ellipsoid 0* and *Point On Ellipsoid 1*. However, the direction from *Point On Ellipsoid 0* to *Point On Ellipsoid 1* can deviate significantly from *Ellipsoid 0*’s outward normal direction at the contact point if an ellipsoid’s aspect ratio deviated significantly from the aspect ratio of a sphere. This leads to non-physical behaviors. To remedy this problem, *Point On Ellipsoid 0* and *Point On Ellipsoid 1* are adjusted to the intersection points between the two ellipsoids and the line having the direction of the outward normal vector and passing through the contact point. The overlap distance *δ*_*N*_ and the direction **u**_*N*_ and magnitude of shoving force were calculated using the adjusted points.

#### 3.3.2 Adhesive Junction Formation

Two cells form a junction if they are compatible and close. Proximity of cells was determined based on the deformed configuration. We increased the principal semi-axis lengths of the two deformed ellipsoids by a parameter *l*_*Ec*_. This expansion represented the extracellular reach of adhesion molecules. A new adhesive junction was formed when the two resulting ellipsoid overlap.

If two cells form a junction, actomyosin contractility in the vicinity of the cell junction pulled the two cells closer [15]. In order to capture this, the MCSM used an equilibrium ratio parameter between two cells with a junction. The two cells were pulled together or pushed apart if they deviated from the equilibrium ratio. To determine the actual junction length and the direction of the force (attraction or repulsion) induced by the junctions, we first found the contact point between the two cells (see Figure2). From this contact point, the model found the nearest points to the two ellipsoids, and these two points became the junction locations for the two cells.

The MCSM computed the junction cross-sectional area using the two ellipsoids used for the closeness test. Two overlapping ellipsoids form a 2D elliptic contact area. The skin base-model assumed that the junction cross sectional area coincided with this contact area scaled by the scale factor (*f*_*C*_ *A*) based on adhesion molecule densities. The junction elastic properties were determined by the junction section area, the junction length, Young modulus, and Poisson ratio assuming a Hookean law. A junction broke if the junction was stretched more than the maximum allowed stretch length from the junction length in the undeformed configuration. Junction directions, cross-sectional areas, and equilibrium ratios were updated in every baseline time step as ellipsoids’ positions, orientations, sizes and junction stretch lengths changed.

#### 3.3.3 Forces

Once the contact point and the penetration depth *δ*_*N*_ was computed, the normal contact Force *F*_*N*_ was computed by the following equations:

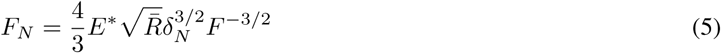

Where

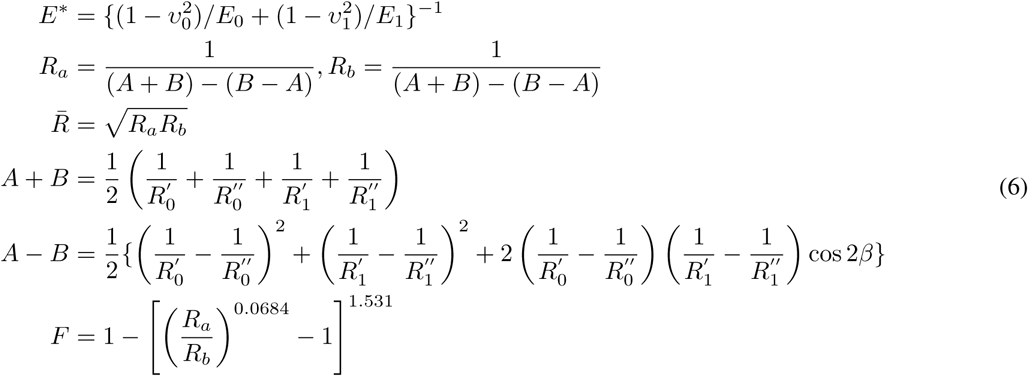

Where *E*_0_ and *E*_1_ were the modulus of elasticity of the ellipsoids, *ν*_0_ and *ν*_1_ were the Poisson ratios; *E*\*was the equivalent modulus of elasticity.; 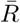 was the effective radius; 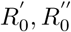 and 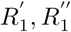 were the principal radii of curvature for the ellipsoids; *β* was the inclined angle [16]; and *F* was the correction factor factor accounting for the non-circularity of the contact area.

We tested our implementation by comparing the computed normal force between two ellipsoids located at different configurations. We expected to reproduce the results of Figure 4 of [17] in which the normal contact force, *F*_*N*_, between two spheres at different separations was computed with Finite elements, and their implementation of computing *F*_*N*_.

**Figure 4:**
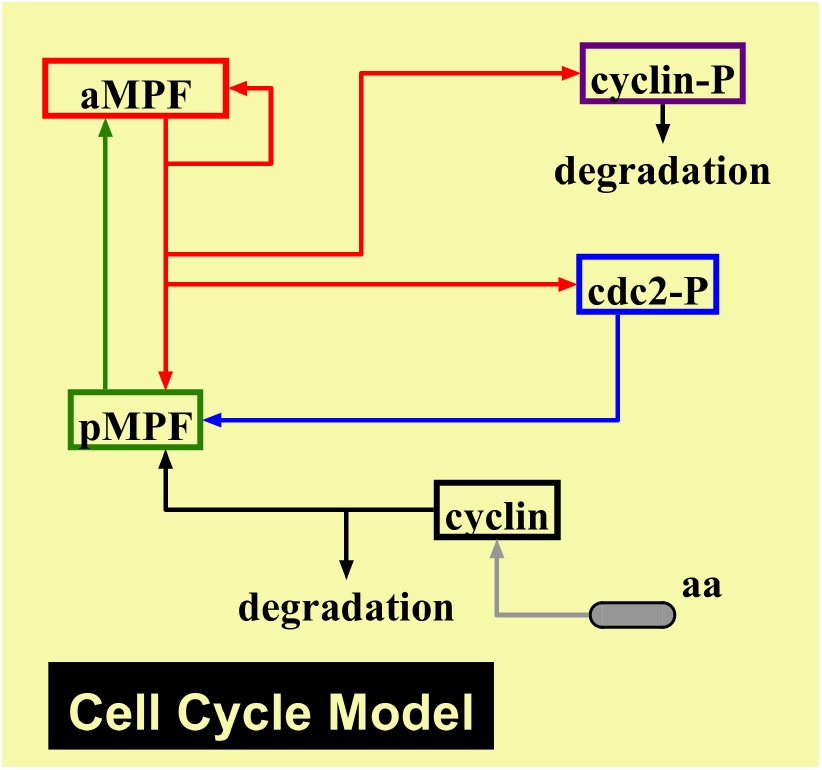
cell-cycle model implemented in the MCSM adapted from [4].

#### 3.3.4 Deformation

Cells could deform from their original (undeformed) shape due to mechanical stresses induced by neighboring cells. The deformation state of an ellipsoid was characterized by the stretch ratios (*λ*_*i*_) along the principal axis:

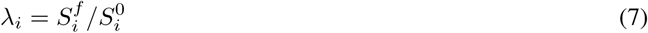

where 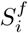 and 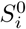 were the lengths of principal semi-axes of the ellipsoid (*i* = *x, y, z*). To find the *λ*_*i*_ values we needed to estimate the Cauchy stresses, which were computed by dividing the normal force by the area in the deformed configuration of the ellipsoid. For simplicity, we used the cross-sectional area of the ellipsoid of the current configuration (before the deformation). Moreover, we only considered forces that were orthogonal to the surface of the ellipsoid. A smoothing factor was applied to the stresses to limit the deformation and reduce the numerical errors of the approximation.

The values of *λ*_*i*_ were then computed from the estimated Cauchy stresses, assuming the cells behaved as a neo-Hookean and incompressible material [18]. It is worth noting that due to smoothing factor the magnitude of the cellular deformations was still in the linear regime for all simulations described here.

## 4 Underlying Models

### 4.1 Agent-Based Cellular Model

The core of our MCSM was an agent-based skin model we previously developed as described in Li et al. [1]. In the original model, we included five different cell types representing different stages of keratinocyte differentiation from stem cells to stratum corneum cells. Here, we included three distinct agent types based on functional differences: proliferating keratinocytes, differentiating keratinocytes and terminally differentiated corneocytes. These were broken down further into sub-types or states as needed.

Proliferating keratinocytes were represented as spheres in their undeformed state 3.1. This model allowed for different states of proliferating keratinocytes, e.g. stem and progenitor or transient amplifying cells as in the original agent-based model [19]. Different division potentials, infinite or finite, as well as division and differentiation rates could be manually set.

We included differentiating keratinocyte sub-types as spinous, granular, and transition. In the spinous state cells were still treated as spheres when undeformed, but by the granular state cells began to polarize and take on a non-spherical resting state (see undeformed in 3.1). They also started growing at a constant rate and increased along the body fixed horizontal direction. Once in the transition state, cell reduced in volume. Biologically, this results from the release of lipids in the extracellular space. Here the volume reduction was preset and we did not explicitly represent intra and extra cellular lipids. Polarization continued in the transition state as the reduction of volume was achieved by reducing the principal semi-axis length only in the body-fixed vertical direction. This may not be the z-axis if the cells have rotated.

Corneocytes were divided into two different sub-types for new and old corneocytes. New corneocytes increased in dry volume growing in the body fixed horizontal directions. In old corneocytes corneodesmosomes, the adhesion molecules degraded and cells underwent desquamation as a result. Corneocytes could change water volume (described in section 4.2.1), and the model assumed that the principal semi-axis length in the body-fixed vertical direction changes due to water volume change.

#### 4.1.1 Agent Cell-Cycle and Proliferation

The current MCSM assumed that proliferating keratinocytes advanced their cell-cycles through four states: G1, G1 checkpoint, G2, and M. Unlike the original model that used a “hard-coded” time, our current version advanced through cell states using the ODE-based cell-cycle model adapted from the Tyson group [4]. This model represented the formation and dissociation of Maturation Promoting Factor (MPF), which is an activated, i.e. phosphorylated, heterodimer of Cyclin and CDC2 4. This system could operate as an excitable oscillator, which represented a typical cell-cycle with metaphase occurring at peak MPF concentration. The model was implemented by a set of five ODE equations that explicitly modeled the individual species, five state variable:

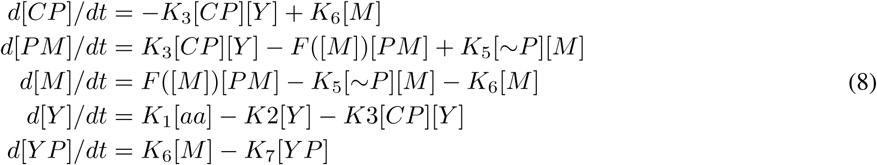

Where [*CP*], [*PM*], [*M*], [*Y*], and [*Y P*] represented the concentration of phosphorylated CDC2, phosphorylated MPF, activated MPF, cyclin, and phosphorylated cyclin, respectively. The concentration [*aa*] and [∼*P*] were held constant. Moreover, it is assumed that CDC2 was immediately phosphorylated so that the total number of CDC2 molecules [*CT*] = [*CP*] + [*PM*] + [*M*]. The parameters were the same as in the original Tyson paper [4], but time was scaled such that the default cycle length was equivalent to the default division time for our proliferating keratinocytes. As in the original Tyson paper, the ODE system could also be simplified into a two state variables, which would be appropriate if only the dynamics, and not the specific individual species, were needed.

The ODE system was suspended during the G1 checkpoint state. From the G1 checkpoint state, a cell advanced to the G2 state only once a randomly chosen number was smaller than a threshold *θ*_*G*1*c*_. G1, G1 checkpoint and mitotic threshold values are tuned assuming that G1 cycle length and G2 cycle length were the same and M cycle length was 10% of the cell-cycle length excluding G1 checkpoint. The threshold was computed as a function of cell “stemness” or division potential, contact inhibition, and TEWL. Specifically, the threshold value was scaled by a factor if the the cell was under contact inhibition; similarly, the threshold value was scaled by three different values corresponding to high, medium, and low TEWL, respectively. Proliferating keratinocytes grew in the G1 and G2 states and divided at the end of the M state.

The traditional view of skin homeostasis includes a central role of slow cycling stem cells, with infinite division potential, giving birth to faster cycling transient amplifying cells. One modeling option would have been to set stem and progenitor cells with different division potentials. However, it is unclear if any explicit distinction between stem and other progenitors is needed to describe normal homeostatic conditions and colony dynamics [20, 21]. For simplicity, we therefore set all basal layer keratinocytes to have infinite division potential while in contact with the basement membrane. Similarly, the default proliferation rates/cell-cycle lengths were the same for all proliferating cells.

The MCSM inherited hard coded probabilities for symmetric vs asymmetric division, where one daughter cell is further along in differentiation, and horizontal vs vertical division. These probabilities could be defined for each proliferating state and tuned to maintain stability of homeostasis and colony dynamics. However, we again used the same parameters for each proliferating state, stem and progenitors. In fact, in our current MCSM, asymmetric and vertical division probabilities were set to zero while differentiation and upward migration result naturally from biological and mechanical interactions.

Ultimately, this implementation had no meaningful difference between any proliferating keratinocyte states. We do emphasize that multiple *modes* of proliferation were possible, but rather than an irreversible state change from stem to progenitor, we considered queues in the local niche, e.g. contact inhibition, as stated above. This provided *interconvertible* modes of proliferation, e.g. expanding and balanced modes [22]. Furthermore, models invoking a strict stem/progenitor hierarchy were indistinguishable from more pliable cell state models for describing colony dynamics [23]. We recognize that work has supported the role of two distinct cell populations, but this distinction becomes more clear in wounding [24, 19]. For our purposes, the added complexity was simply not needed; however, the code to introduce a distinction exists and can be activated if needed in the future.

#### 4.1.2 Agent Delamination, Differentiation and Differential Adhesion

Previous studies have shown that the basal layer of skin is in a jammed solid-like state and delamination events occur in response to neighboring division events, which increased the local density and stresses [25]. In this MCSM implementation, basal cells initiated differentiation in response to these mechanical forces rather than idiopathic probabilities of symmetric or asymmetric division. Specifically, differentiation began either upon delamination, when a cell was mechanically forced from the basement membrane, or when the cell was vertically elongated due to lateral compression. The elongation was a natural result of cells, modeled as elastic bodies, subjected to increases in local density. Delamination was driven mechanically and facilitated by changes in the adhesion complexes expressed of the surface of dividing cells [25].

From this point, differentiation through the spinous, granular, and transition states took place as a function of time and calcium gradient. It is important to note that the calcium profile was static; although, models of calcium transport dynamics are available for future implementation [26]. Differentiated keratinocytes differentiated to corneocytes passing the transition state. Corneocytes became detached once they lost all cell-cell junctions.

Differentiated keratinocytes polarized once they entered the granular state and begin to flatten out in the default undeformed state after losing lipids, which happened as part of the predetermined differentiation process. The vertical direction was re-evaluated as cells may rotate after reaching this state and their body-fixed z direction may not be the desired vertical direction. The new ideal vertical direction was reevaluated by finding the body-fixed axis closest to the global z axis. Cells also gradually adjusted their orientation to match the selected body-fixed axis to the global z direction.

As cells delaminated, differentiated and migrated upwards they displayed different adhesion complexes on the surface. This resulted in differential adhesion forces and cell-cell contact areas (described in 3.3.2) as cells of the same type tended to form stronger interactions. Differential adhesion can facilitate self-organization in developing biological systems [27] and may play a role in maintaining robust, well-defined skin layers in vivo. As in Du et al. 2018 [10], we also noted a sharpening of the layer boundaries in our simulations when differential adhesion was implemented (data not shown).

### 4.2 Transport Models

Water and molecular transport were modeled using preexisting, continuous transport models that employed one-dimensional PDEs. We leveraged Biocellion’s built-in PDE capabilities to reconstruct and solve these systems in our model.

As described in section 2.2, we used grid spacing of 50*µm* with 16 sub-grids in each dimension. PDE inputs were constant for all sub-grids in a grid box except in the neighborhood of the stratum corneum where we applied the resolution of 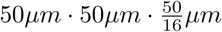. We partitioned the cubical grid boxes near the stratum corneum into thin but wide slabs because parameters varied faster in the z-direction.

#### 4.2.1 Cell Swelling and Water Transport

Water transport and TransEpidermal Water Loss (TEWL) was modeled after Li et al. 2015 [3]. They assumed the stratum corneum to be a rate-limiting homogeneous slab. A PDE system was then set up over this slab with a constant bottom source term. Static diffusion coefficients were calculated based on the physical and biological properties. Finally, they applied a moving upper boundary condition to represent stratum corneum swelling as corneocytes increased water volume.

In our system the stratum corneum was treated as multiple slabs and water was stored in discrete cells that overlapped these slabs. However, the water volume in each slab was still considered to be homogeneous by using a cell overlap weighted average of the cellular water volume at the time of each grid time step. The diffusion coefficients within slabs were defined locally and dynamically as a function of water volume, and the stratum corneum boundaries were defined locally and dynamically based on the location of the corneocytes. Corneocytes explicitly absorbed or released water based on changing local water volume and preset rates. The principal semi-axis length in the body-fixed vertical direction changed as a result of cellular water volume changes. The swelling of individual cells then gave rise to the macroscopic swelling of the epidermis without the need to derive complex boundary conditions. Other cells were assumed to have a constant water volume and act as an infinite source. This could be easily changed by reprogramming the properties of different cell types and defining other boundary conditions, which Biocellion allows for on the faces of any grid box.

For the outermost slabs, we computed TEWL as a function of relative humidity, which provided influx, and the water ratio of the slab, which provided outflux. The resulting TEWL could then be a source or sink term for water transport. A slab was considered to be outermost if there was no other slab having overlapping agents within 50*µm* above the slab. As a simplification there was no transport gradient of water vapor into the bulk atmosphere and the relative humidity was statically defined at the surface.

Taken together, this defined the location of the PDE grids/ sub-grids, the water concentration in each sub-grid, the resulting diffusion coefficients and the source/sink terms. This provided a similar PDE system to the original model, but location, thickness and boundary conditions of the stratum corneum were determined by a biologically dynamic morphology. We also solve the PDE in three-dimensions, but the system at hand had a high degree of axial and radial symmetry in the x/y plane, so this advantage was admittedly minimal.

We note that such transport equations are intended for infinitely dilute solutes and that water, in this case, did not meet this definition. Furthermore, this system assumed the stratum corneum is a mostly static, homogeneous slab while in fact it is a complex packed bed of cells and lipids. Local geometries of the skin can change and large interstitial spaces can open as a result of certain perturbations. Water transport is further complicated by passages directly between cells via cell-cell junctions (e.g. gap junctions). There may even be a role for adhesion complexes such as desmosomal junctions. In addition, water can also be present in bound and unbound states, regulated in part by natural moisturizing factors within cells. All of these biological components can dynamically respond to perturbations.

Importantly, the imposed cubic grid on the extracellular space could be used to calculate local transport properties based on the resident cells, free volume and its material composition. Moreover, the agent-based modeling framework allows us to explicitly define mechanical and biological processes at the level of discrete cells. Forming these discrete systems is far simpler then deriving and re-deriving continuous governing equations (which may not even be possible in some cases) every time a new rule considered. However, the diffusion based transport of the underlying sub-model was shown to be a reasonable approximation under homeostasis or when changes occur faster then typical biological processes. This is an example of how a sub-model defined by continuous PDEs can be directly incorporated into the whole system model while still providing clear opportunities for future refinement.

#### 4.2.2 Skin Penetration

Similar to TEWL, skin penetration by small molecules was modeled after a preexisting, continuous transport model that employed one-dimensional PDEs [2]. This system considered the stratum corneum, viable epidermis and dermis as three distinct homogeneous layers with fixed boundaries. Diffusion coefficients were calculated as a function of chemical properties and layer specific biological properties. They were pre-calculated for each layer and were time-invariant. Our model also defined the layer specific diffusion coefficients prior to simulation using the same math; however, we did not use predefined layer boundaries. Instead, we defined local diffusion coefficients by volume-averaging the coefficient for each overlapping agent, which corresponded to one of the three layers. This allowed the layers an opportunity change with a biologically dynamic morphology.

Flux into the outermost slabs, and therefore the source terms for the PDE, were computed as a function of the predefined external species concentration and the concentration of the slab. We assumed an infinite, time-invariant reservoir (i.e. constant external concentration) for the external source of the molecule species. We also included a constant molecular decay rate. Although an absorbing boundary layer could be explicitly defined as the dermal sink term, we simply used a large dermal compartment. The resident agents in this area are static and the PDE boxes are larger than those in the epidermis.

As with TEWL there are many opportunities to define parameters locally, as opposed to large homogeneous layers. However, for this proof-of-concept, we maintained a simple description of the system as to recapitulate the original transport model.

## 5 Model Simulations

In this section we present several simulations to verify the proper implementation of the sub-models and to demonstrate how the MCSM can be used to study environmental, chemical and physical perturbations.

The MCSM represents an extensive body of model code. While the framework provided handling of variables, partitioning across nodes and numerical solvers, the model code implemented biological, chemical and mechanical rules; PDE/ODE setup; and computational strategies to solve for the mechanics. In all there were thousands of lines to represent this specific model. This complexity made it all the more necessary to confirm model implementation and function, but at the same time it made this process difficult. In this case, we opted for a typical software approach by implementing the concept of unit tests to ensure expected function of critical model aspects. We have already implemented unit tests relating to the physics (Fig 3).

### 5.1 Verified cell-cycle Sub-Model

We verified the proper implementation of the cdc2-cyclin cell-cycle sub-model [4]. We sampled individual progenitor cells in the basal layer and tracked the concentration of their intracellular molecular species. We demonstrated the same oscillatory pattern as in the original Tyson paper. For example, the dynamic behavior of both total cyclin and active MPF relative to total cdc2 (Fig 5) displayed the same limit cycle oscillations as in Fig 3 of [4].

**Figure 5:**
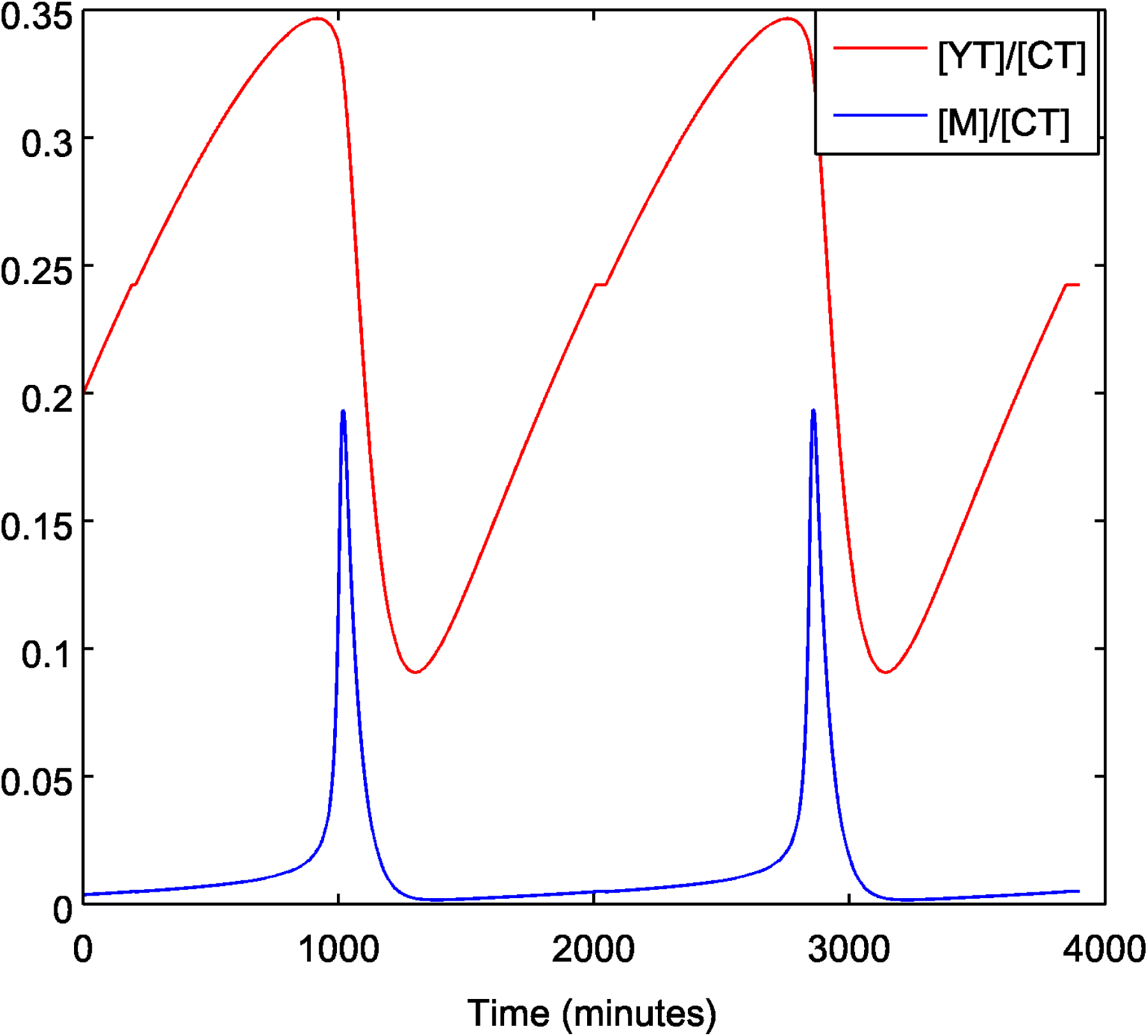
Dynamics of the of the cdc2-cyclin cell-cycle sub-model in our MCSM to be compared to Fig 3A in [4]. Total cyclin ([YT] = [Y] + [YP] + [pM] + [M]) and active MPF [Ml relative to total cdc2 ([CT] = [C2] + [CP] + [pM] + [MI) for the the differential equations 8. Dynamic behavior is consistent with limit cycle oscillations.

### 5.2 Barrier Formation and Homeostasis

Our first test of model integration was to confirm the MCSM’s ability to form a stratified barrier and maintain a stable homeostasis. We seeded 80 cells onto a mostly static basement membrane that consisted of several small (1*µ*m radius) agents in the shape of rete ridges [28]. These ridges were created by placing basement membrane agents in an oscillatory pattern with a hill height of 5*µ*m, valley depth of 50*µ*m and a period of 100*µ*m. Hills and valleys were described by negative and positive exponential functions, which define a steeper slope than traditional trigonometry functions. The only function of the basement membrane agents is to shove or adhere to proliferating keratinocytes in the basal layer -the agents are stationary and do not move despite applied forces. The simulation was run in dry (relative humidity of 0.05) conditions that were arbitrarily chosen.

The simulation was run out for 60 days. The total number of cells came to an approximate steady-state just after 20 days (Fig 6), consistent with the epidermal turnover in young adults [29]. Looking at the agents that make up different epidermal layers indicated more complex dynamics (Fig 6 Right). The stratum corneium, indicated by number of corneiocytes (cc), did show modest oscillations out to 50 days. Interestingly, we observed overshoot of all layers with the most notable in the cells making up the spinous layer. This suggested a significant role for negative feedback in reaching homeostasis. The most predominant form of negative feedback was the relationship between proliferation and TEWL, as suggested in the original model by Li et al. [1]. TEWL is inhibited by barrier thickness so the forming barrier would slow itself down with a delay propagating through the layers.

**Figure 6:**
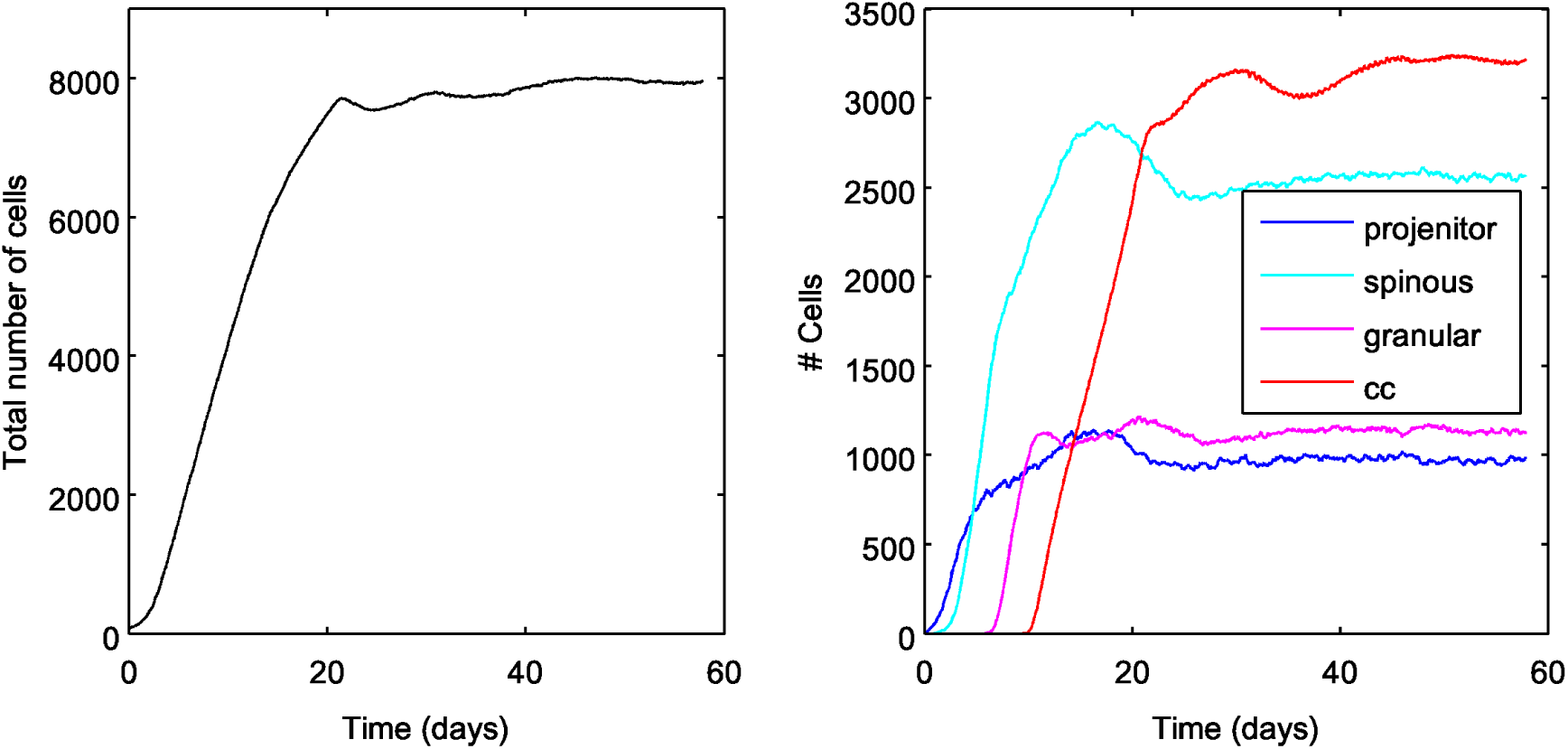
MCSM simulation from low density seeding of the basal layer to 60 days. Left shows all cellular agents in the system and right shows agents comprising individual layers. There is noticeable overshoot, particularly in the spinous layer, which indicated strong negative feedback. Despite some minor fluctuation out to 50 days, the overall epidermal system is near homeostasis prior to 25 days, similar to the clinical rate of epidermal turnover.

### 5.2.1 Environmental: Impact of Relative Humidity (RH)

The cell swelling and water transport sub-model is sensitive to the external RH. We simulated different RH levels and ran the simulation to steady state. We then recorded the average TEWL and stratum corneum thickness (Fig 7. We saw a similar relationship as observed in the original TEWL and cell swelling paper [3]. There is an exponential increases in SC thickness with increasing RH accompanied with rapidly decreasing TEWL. The TEWL decreases due to both the increased distance water must pass thought the SC and the reduced gradient between the sink to the external environment and the source from within the skin.

**Figure 7:**
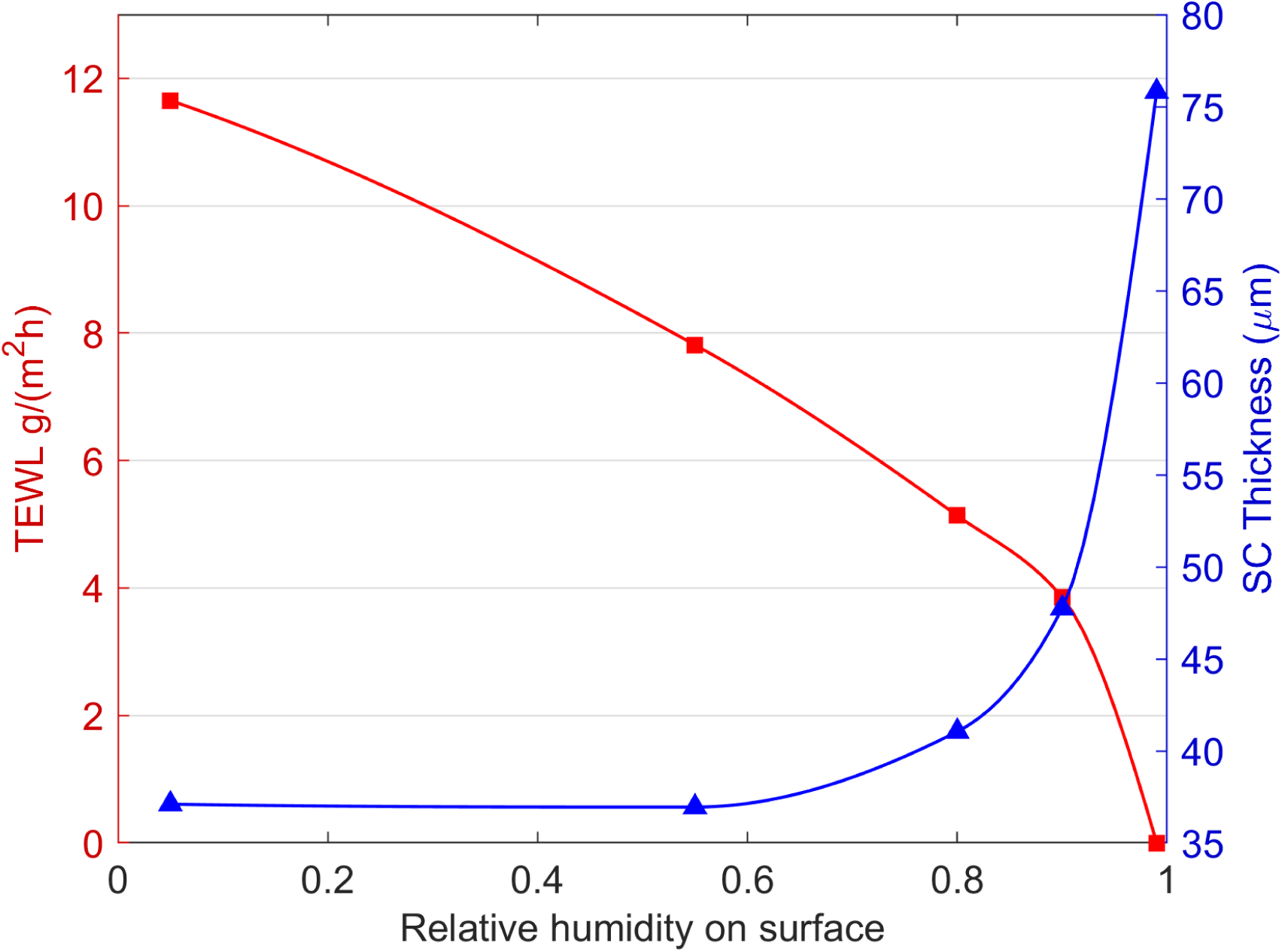
Stead-state values of stratum corneum thickness (blue) and transepidermal water loss (red) as a funciton of relative humidity. To be compared to Fig 3 in [3]. The similarity to the original cell swelling and water transport validated the implimnetation of this sub-model.

#### 5.2.2 Chemical: Topical Treatment of cell-cycle Inhibitor

Next we considered chemical perturbations, which typically occur from topical application of a material that contains chemical agents that can penetrate the stratum corneum and interact with cells in the viable epidermis. As suggested by the description, this will primarily depend on the interplay between the skin penetration of a chemical species and cellular agent-based sub-models of the MCSM. If a biological mechanisms is known, an intracellular ODE reaction system can bridge the chemistry to the affected biological processes. To demonstrate this we considered an established cell-cycle inhibitor, specifically CDK1 (AKA CDC2) inhibitor CAS 220749-41-7. The sub-model of inhibitor transport is described in section 4.2.2, and the values of the parameters are shown in “cell-cycle inhibitor”.

To model the effect of cell-cycle inhibitor we extended the five variable model of cell-cycle adding two extra variables that represent the intra-cellular concentration of the Inhibitor ([*I*_*int*_]) and complex composed of the Inhibitor and the active PMF([*MI*]). The seven state variable system is basically the same as the original five variable system, but includes an additional interactions by which the inhibitor decreases the concentration of CDC2. The following equations are added or changed from the five variable cell-cycle model:

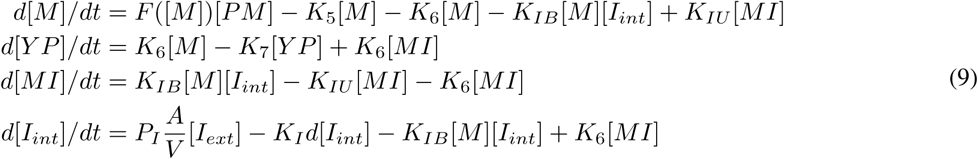

Where *A* and *V* are the area and volume of a cell, respectively. *I*_[_*ext*] is the extracellular concentration of the inhibitor which is computed by the penetration model as mentioned above. *K*_*IB*_, *K*_*IU*_, and *K*_*I*_ are estimated considering that at half maximal inhibitory concentration (EC50), 50% of *M* is bound to the inhibitor, then at steady state [*I*_*EC*50_] = *K*_*IU*_ */K*_*IB*_. Assuming that [*MI*] dissociate at the same rate than [*M*] dissociation, then *K*_*IU*_ = *K*_6_. The EC50 of inhibitor of CDC2 is 5.8*µ*M and *K*_*IB*_ = 5.8*/K*_*IU*_

Figure 8 shows how skin properties changed after topical application of the cell-cycle inhibitor. In the first 15 days, most of the inhibitor was only able to diffuse inside the stratum corneum, see Figure 8A. This was due to the high diffusion coefficient of this layer. Yet, there was enough inhibitor penetration to impact proliferation and progenitor cell number after days of exposure (Fig 8B). However, barrier thickness and TEWL were not affected in this period (Fig 8 C).

**Figure 8:**
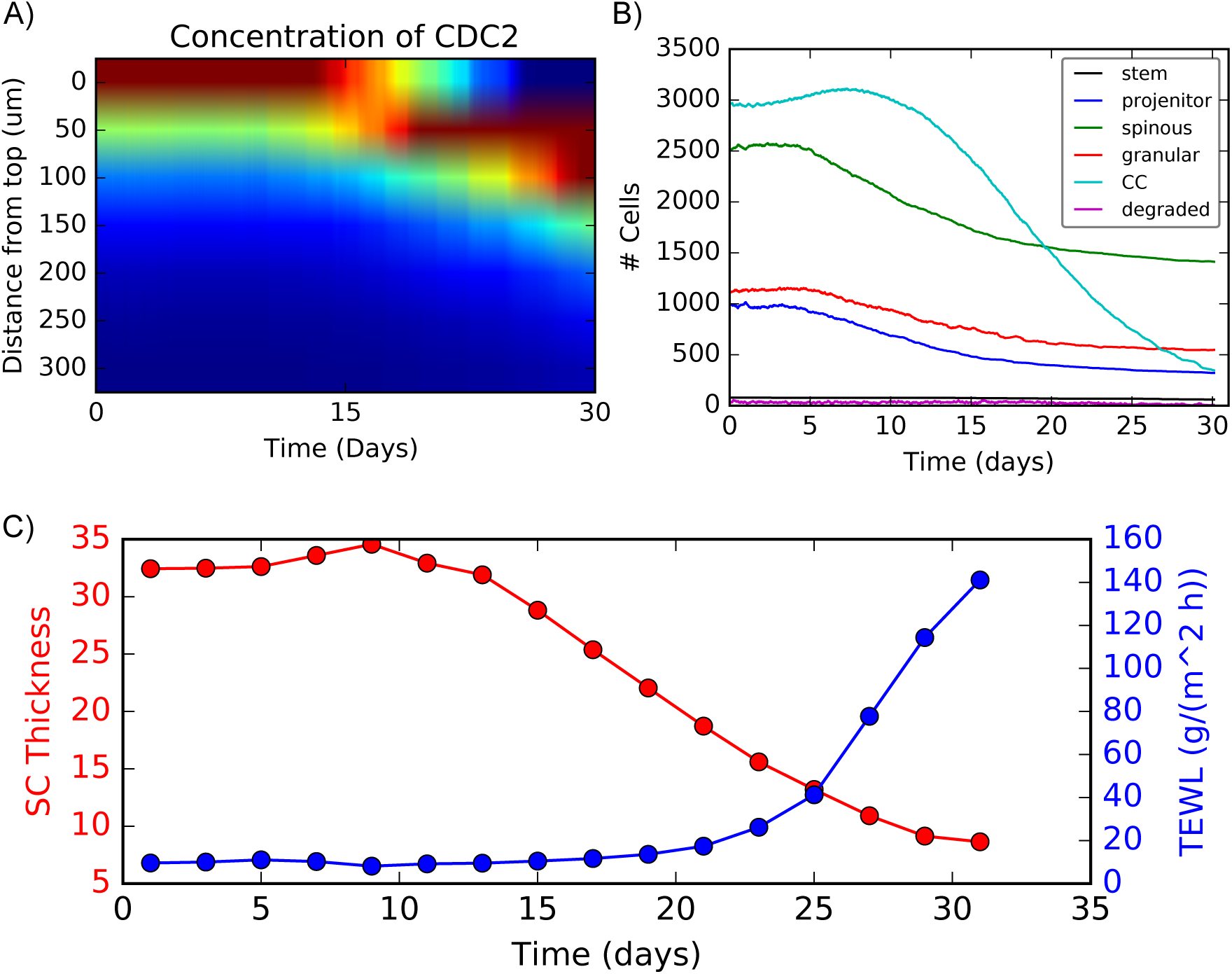
Inhibition experiment. A: Heat map of average concentration of inhibitor; red and blue colors indicate high and low concentration respectively. B: Population change after topical application of the inhibitor. C: temporal change in SC thickness and TEWL after topical application of inhibitor.

As this process continued, fewer differentiated cells were available to maintain the upper layers. Eventually the stratum corneum, which was constantly being sluffed off, was not sufficiently replenished with corneocytes leading to barrier disruption, about 10 days. This corresponded with a slow rise in TEWL (Fig 8 C). After approximately 15 days, the inhibitor rapidly diffused through the viable epidermis further inhibiting proliferation and rapidly reducing the population of cells in the lower layers. After 20 days TEWL began to rise rapidly, demonstrating the overall loss of barrier function.

#### 5.2.3 Mechanical: Micro-indentation

The mechanical properties of the skin are of importance for various cosmetic and clinical applications. The MCSM model can be used to perform virtual mechanical experiments commonly used to estimate elastic properties of different materials. We performed uniaxial indentation experiments, simulating the experiments described in [30], that consisted of compressing a segment of skin by using an spherical probe.

The indentation experiment assumed that simulation begins from a fully developed skin sample. Before applying stress using a spherical indenter, we cut a rectangular sample from the developed skin and setup periodic boundary conditions in the x-y directions and a hard wall boundary conditions in the z direction. Cell division, cell death, cell differentiation, and transport of water were not modeled as they occurred at time scales much larger than the indentation experiments (which was simulated in seconds).

The height of the spherical probe was initialized in such a way that there was no overlap between the probe and the skin segment. The probe was then move downwards slowly with constant velocity. At every baseline time step we computed the average force on the spherical probe due to mechanical indenter interactions with cells of the skin segment. Figure 9 shows the force-displacement curves obtained by the computational indentation experiments. Stiffness was characterized by the slope of the curve at the initial (linear) part of the force-strain experiments. It is worth nothing the non-linearity of the force-displacement curve generated by the indentation experiment. A linear regime is followed by constant force movement of the indenter.

**Figure 9:**
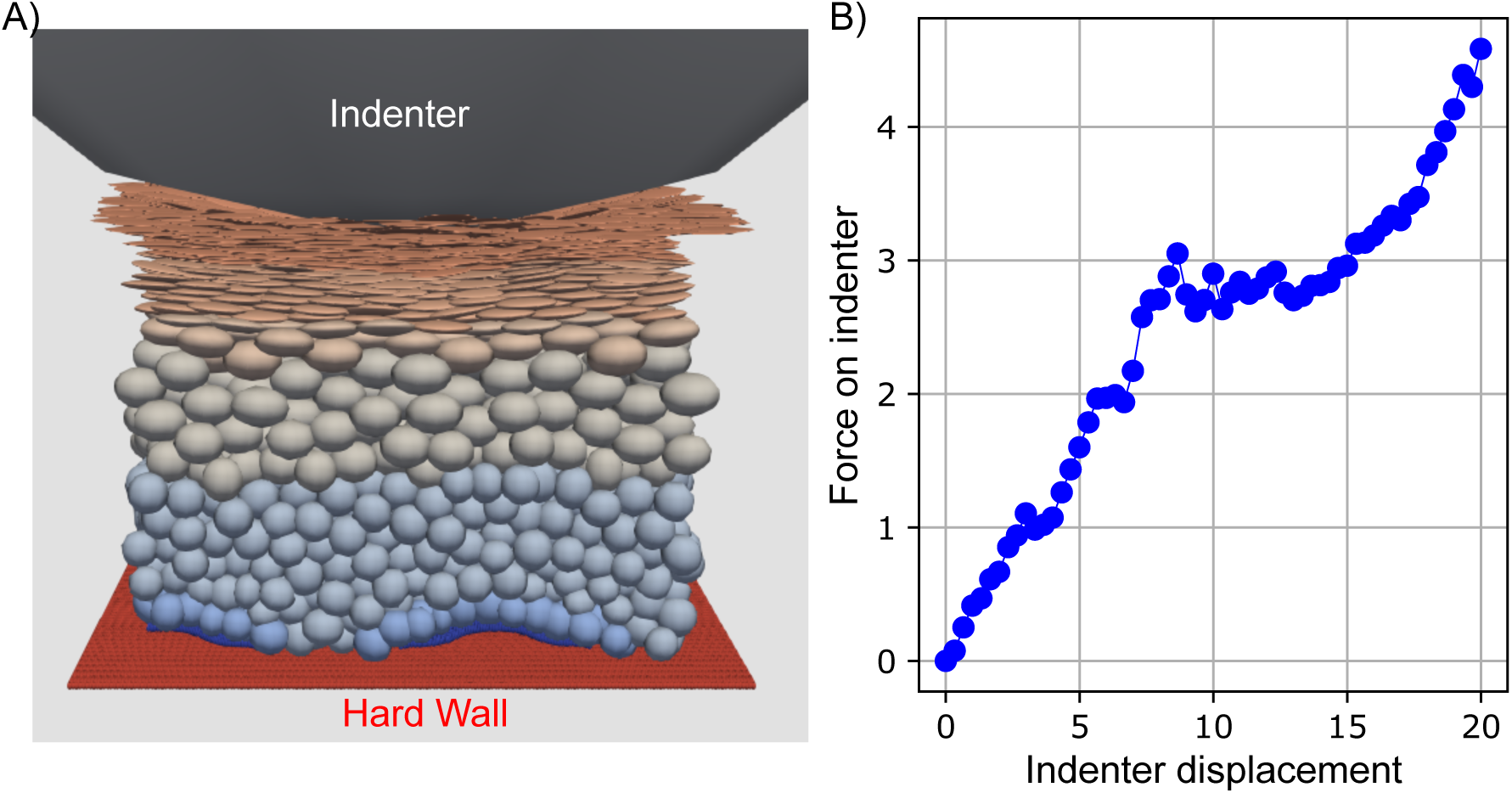
Indentation experiment. A: Snapshot of the indentation experiment. B: Force-displacement curve from the indentation experiment. The force on indenter is computed by summing the vertical component of the forces between the indenter and cells.

## 6 Discussion

Taken together, our multiscale MultiCellular Skin Model (MCSM) was able to recapitulate results from the original skin models and, importantly, produced the expected results for more complex conditions requiring model integration. For example, the cell-cycle oscillations (Fig 5) as well as the impact of RH on thickness and TEWL (Fig 7) were meant to be narrow recapitulations of the corresponding original publications.

Alternatively, the formation of the skin barrier was a result of integrating rules from proliferation and cell-cycle to delamination and differentiation. The control of these processes created a balance that lead to a stable homeostasis of skin morphology (Fig 6). Interestingly, we achieved this balance with no distinction between stem and progenitor proliferating states (section 4.1.1. Furthermore, we did not impose idiopathic, predetermined probabilities for asymmetric or vertical division to manually control differentiation and vertical migration. Instead, the integration of biology and mechanics, absent from the original model, was sufficient. This was also consistent with experimental observations [25].

Perturbations that lead to an imbalance in proliferation and differentiation can result in the destruction of the skin barrier. In order to simulate this, we modeled skin treated with a cell-cycle inhibitor (Fig 8). Importantly, this simulation brought together all of the underlying sub-models and demonstrated negative feedback as a result. Penetration of the inhibitor into the skin was described by the skin penetration sub-model (section 4.2.2), which determined the bioavailability to cells in the basal layer. This in turn determined the concentration of inhibitor available for binding the CDC2 complex, which slowed the Tyson cell-cycle model (section 4.1.1). Ultimately, the skin barrier weakened as dictated by the biology and mechanics of the agent based model (section 4.1). The weaker barrier allowed more penetration of the cell cycle inhibitor-negative feedback. Eventually the barrier collapsed and the water transport model (section 4.2.1) predicted a precipitous rise in TEWL. Linking the material application to mechanism and finally a clinical measure of barrier required each sub-model, and the important role of negative feedback could only be achieved by an integrated system as opposed to a serial workflow.

It is worth noting that no parameter tuning was done specifically for this experiment. The transport properties were derived from the chemical properties of the inhibitor. The interaction of the inhibitor and its target was estimated from the EC50 value. The remaining cell-cycle parameters were unchanged. The ultimate destruction of the barrier was then a result of the same rules and parameters used in simulating barrier formation.

Finally, the results of the mechanical perturbation demonstrated an emergent behavior. The mechanical properties of the whole tissue were a manifestation of the underlying mechanics of the cellular agents. Here, we modeled individual cells as well as cell-cell junctions as elastic bodies with deformation within the linear range (section 3.3.4). Using Hooke’s law to model the whole tissue as a single slab, would have resulted in a force-displacement curve similar to an individual cell (Fig 3). However, our cell-centric model produce a highly non-linear results (Fig 9) including incomplete elastic recovery. In this case agents can be reorganized under a certain stress and perhaps get stuck in a new local free energy minima. This example demonstrated how an unexpected non-linear system behavior can emerge from the more simplistic behavior of the underlying components.

### 6.1 Limitations

As with any model, we have simplifying assumptions, constraints and other limitations. Here, we attempted to capture only the most basic elements of skin barrier function -the biology and balance of proliferation and differentiation, cellular mechanics and the physics of mass transport. Integrating these elements allowed us to capture basic features of barrier formation and homeostasis. It also allowed us to demonstrate the possibilities of modeling skin response to various perturbations. While there are many limitations of this simplistic view, as delineated below, we suggest that further model development be done only with a specific condition and output in mind.

There are significant limitations on the molecular biology. Molecular biology, in particular, tends to encourage complexity in order to capture known mechanistic details. This can introduce significant challenges such as parametric complexity (discussed below). Here we purposefully limited ourselves to a single, highly simplistic, biomolecular interaction network for eukaryotic cell-cycle. This was included only to link the molecular mechanism of our topical inhibitor to the cellular function of proliferation. The only other explicit biomolecular level detail was a limited set of adhesion molecules that determined cell-cell junction strength. The model could not make use of, nor output, any other biomolecular information. Of course, additional details could be integrated and the cell-cycle network could be used as a template.

Notable limitations on biological function included no model for apoptosis, active cellular migration or inflammatory response, as well as a limited differentiation model. In the original model an asymmetric division hypothesis, implemented as a set probability, was used to determine cell fate post division. Stems cells could have different set probabilities. In this model there was no meaningful difference between stem cells and other proliferating keratinocytes. Both are exposed to the same mechanical mechanism for delamination and subsequent differentiation. Once differentiation is initiated it continues merely as a function of time and calcium concentration. These limited mechanisms were sufficient for our purposes. More dramatic perturbations, like deep wounding or long-term aging, are expected to require a more detailed descriptions of stems cells, polarization and the overall differentiation process.

The only explicit small molecular effect on biology was between calcium and differentiation state. Importantly, pH was not considered in the model. pH has a broad impact on biological function and is particularly important in skin, which has an acidic external surface. We implicitly assumed that pH is not modulated under the conditions considered here, or at least not in a way that impacted our reported outputs. However, there are conditions where pH plays a major role and would need to be included. One example is the maturation of preterm or infant skin in which the external surface pH changes from neutral to acidic in the days following birth.

Mechanics are limited to fully elastic bodies. As of yet, we have had no need to include a complete viscoelastic model. The geometry of the bodies themselves were limited to small rods for junctions and ellipsoids for cells. The use of simple ellipsoids worked well in the basal layer where cells undergo mechanical stress and become columnar. It was also reasonable for the stratum corneum as nearly flat discs could be formed and through cell-cell junctions these can form sheets. Both of these behaviors are characteristic of skin morphology. However, ellipsoids provide a poor description of the middle skin layers. Here cells are tightly packed and deform into complex shapes that minimize the free volume between cells. We cannot capture this morphology in our current framework. A more appropriate description for tightly packed tissues may be vertex or boundary modeling. The cellular Potts model, for example, could capture this feature.

Mass transport limitations arise from how transport properties were defined. We limited the model to pre-calculated transport properties that differed only by skin layer. We did not take into account local compositions or geometries. We did not account for vacuoles which can form under certain conditions. We did not account for lipid composition. The calculation of the transport properties implicitly account for lipids since the parameters were measured in normal skin, but there was no way to account for changes in lipid composition in the current model. Lipids could be accounted for either by making them properties of the sub grids that can be altered by cells within the grid or they could be defined as species within the context of the PDE and be secreted by cells. For the latter, advection would need to be included to capture the bulk movement due to upward migration of cells. Other limitations, challenges and opportunities were discussed in sections 4.2.1 and 4.2.2.

### 6.2 Parametric Complexity

Parametric complexity is often the most concerning element in modeling. When modeling biological processes, we are often met with a trade-off between including the high complexity of biological systems versus a practical statistical issue of a high dimensional parameter space. Over simplification upsets the biologists and over parametrization worries the practitioners of the physical scientists. This is often the case in ODE-driven compartment models of cellular processes or pathway modeling. It may seem natural to include all the known (or suspected) players (proteins and other biomolecular species) and their interactions. However, this often produces high-dimensional systems with hundreds and sometimes thousands of mostly unknown parameters that are too difficult to measure directly. Combined with sever limitations on biological data, these systems are massively undetermined. There are methods, sometimes quite insightful, for mitigating some of this trade-off; however, we will not delineate these here, and we simply suggest that it is still a major challenge in the field.

Our model included 345 total parameters, or more precisely preset static variables. Despite the number of parameters, we maintain that our model did not suffer from the same degree of parametric complexity. This was proven out at a practical level as we found no need for an optimization algorithm and only minimal manual tuning to produce all the simulations provided here. There were several reasons for this. First, there were far fewer free parameters that represented the functional biology/physics in a meaningful way. Some parameters represented details for simulation implementation, numerical solvers and error conditions; some were placeholders for future expansion or increased detail when needed; some were set as functions of other parameters; and some were duplicated over all agent types and changed only if a distinction was needed. Additionally, some sub-sets of parameters were only used under specific simulation conditions. Secondly, several parameters were reasonably constrained or explicitly set based on the scientific literature. Finally, we have attempted to choose the appropriate scale when modeling each process to facilitate the use of data that is more easily obtainable (in replace of detailed mechanistic knowledge). Of course, if such detail is available, the flexibility of our framework would allow for implementation.

Cell proliferation is a good example of capturing a process at the right scale. Our proliferation model did include details at the molecular level. However, the parameters were constrained by the literature, and we also give the user options to effectively course grain over the mechanistic detail when it is not needed. The five molecular variables could be analytically reduced to two system variables. In addition, we allowed the user to set the default cell-cycle length. If the mechanism is known, like in our inhibitor example, the molecular pathway can be expanded and rate constants can be adjusted. If the mechanism is unknown, the user can modify the cell-cycle length directly. Recall that the dynamics of the cell-cycle pathway were scaled by default cell-cycle length (Section 4.1.1). Measuring a parameter for the rate of proliferation is far easier than measuring reaction rate constants, and simple *in vitro* systems could determine a functional relationship between concentration and proliferation rate. Knowledge of mechanistic detail can be offset by direct experimental data and vice versa, and this ability is built into our modeling framework and implementation.

### 6.3 Rising Concerns with Mounting Complexity

We have already pointed out the difficulties of validating the *implementation* of the code much less the underlying biological and physical principles. But in fact, the issue is deeper still. With thousands of lines, the model code is approaching the limit of human interpretability. It should be observed from this document that providing a description of the model that is both precise enough to maintain scientific rigour yet easily digestible was quite difficult. Obviously, we are well beyond elegant analytical solutions and the deep insights that arise, but it is becoming difficult to simply conceptualize the model as a whole. While the description of the multiple sub-models can be daunting in its self, understanding the integration at the needed precision is the most difficult part. The primary issue at hand is the use of fundamentally distinct physical concepts, mathematical frameworks and numerical solutions for a single system.

This multifaceted modeling approach is in contrast to something like a compartment, process or pathway model that may limit its implementation to ODEs. An ODE reaction system may be meaningfully abstracted into a network diagram. If the field agrees to a small set of descriptors for the reaction details (e.g. mass action kinetics) then the diagram can be a precise description of the model. Even if thousands of interactions make such a diagram difficult to interpret as a whole, the precise and systematic abstraction makes it easy to interpret individual processes or modules. Importantly, it also makes sharing the model significantly easier.

However, when modeling tissues at cellular resolution, there is no agreement in the field as to what specific physical or biological concepts should be included and to what level of detail should they be described at. There is no agreement as to level of flexibility a framework should have, and there is certainly no agreement on an implementation strategy. We argue the need to identify a set of ‘good’ models that capture the core physical and biological concepts before we can work on establishing a common ontology or other abstractions as well as a common set of implementation methods in which the details can be easily communicated. By ‘good’ we mean models that the field can agree demonstrate *significant* utility outside of an academic context. We argue that no such set of inter-subjectively useful models in the field of multiscale multicellular tissue modeling exist in the published literature at this time. To this end, future work will be for the application of this model to meaningful clinical and industrial purposes.

## References

[1] X. Li, A. K. Upadhyay, A. J. Bullock, T. Dicolandrea, J. Xu, R. L. Binder, M. K. Robinson, D. R. Finlay, K. J. Mills, C. C. Bascom, C. K. Kelling, R. J. Isfort, J. W. Haycock, S. MacNeil, and R. H. Smallwood. Skin stem cell hypotheses and long term clone survival–explored using agent-based modelling. Sci Rep, 3:1904, 2013.

[2] Y. Dancik, M. A. Miller, J. Jaworska, and G. B. Kasting. Design and performance of a spreadsheet-based model for estimating bioavailability of chemicals from dermal exposure. Adv Drug Deliv Rev, 65(2):221–36, 2013. Dancik, Yuri Miller, Matthew A Jaworska, Joanna Kasting, Gerald B eng R01 OH007529/OH/NIOSH CDC HHS/Research Support, N.I.H., Extramural Research Support, Non-U.S. Gov’t Review Netherlands 2012/01/31 06:00 Adv Drug Deliv Rev. 2013 Feb;65(2):221–36. doi: 10.1016/j.addr.2012.01.006. Epub 2012 Jan 23.

[3] Xin Li, Robert Johnson, Ben Weinstein, Elizabeth Wilder, Ed Smith, and Gerald B. Kasting. Dynamics of water transport and swelling in human stratum corneum. Chemical Engineering Science, 138:164–172 Elsevier.

[4] J. J. Tyson. Modeling the cell division cycle: cdc2 and cyclin interactions. Proc Natl Acad Sci U S A, 88(16):7328–32, 1991.

[5] Tyson, J J eng GM-36809/GM/NIGMS NIH HHS/Research Support, U.S. Gov’t, Non-P.H.S. Research Support, U.S. Gov’t, P.H.S. 1991/08/15 Proc Natl Acad Sci U S A. 1991 Aug 15;88(16):7328-32.

[6] Seunghwa Kang, Simon Kahan, Jason McDermott, Nicholas Flann, and Ilya Shmulevich. Biocellion: accelerating computer simulation of multicellular biological system models. Bioinformatics, 2014.

[7] Ji Won Oh, Tsai-Ching Hsi, Christian Fernando Guerrero-Juarez, Raul Ramos, and Maksim V Plikus. Organotypic skin culture. The Journal of investigative dermatology, 133(11):e14, 2013.

[8] Gavin Maxwell and Cameron Mackay. Application of a systems biology approach to skin allergy risk assessment. Altern. Lab. Anim., 36(5):521–556, November 2008.

[9] Maria F Leyva-Mendivil, Anton Page, Neil W Bressloff, and Georges Limbert. A mechanistic insight into the mechanical role of the stratum corneum during stretching and compression of the skin. J. Mech. Behav. Biomed. Mater., 49:197–219, September 2015.

[10] Georges Limbert. Mathematical and computational modelling of skin biophysics: a review. Proc. Math. Phys. Eng. Sci., 473(2203):20170257, July 2017.

[11] H. Du, Y. Wang, D. Haensel, B. Lee, X. Dai, and Q. Nie. Multiscale modeling of layer formation in epidermis. PLoS Comput Biol, 14(2):e1006006, 2018. Du, Huijing Wang, Yangyang Haensel, Daniel Lee, Briana Dai, Xing Nie, Qing eng R01 AR068074/AR/NIAMS NIH HHS/ R01 GM107264/GM/NIGMS NIH HHS/ R01 NS095355/NS/NINDS NIH HHS/ R56 AR064532/AR/NIAMS NIH HHS/ Research Support, N.I.H., Extramural 2018/02/27 06:00 PLoS Comput Biol. 2018 Feb 26;14(2):e1006006. doi: 10.1371/journal.pcbi.1006006. eCollection 2018 Feb.

[12] Thomas Sütterlin and Niels Grabe. Graphical Multi-Scale modeling of epidermal homeostasis with EPISIM. In Computational Biophysics of the Skin, pages 436–475. Pan Stanford, 2016. Thomas Sütterlin, Erika Tsingos, Jalil Bensaci, Georgios N Stamatas, and Niels Grabe. A 3D self-organizing multicellular epidermis model of barrier formation and hydration with realistic cell morphology based on EPISIM. Sci. Rep., 7:43472, March 2017.

[13] Xiaoshan Lin and Tang-Tat Ng. Contact detection algorithms for three-dimensional ellipsoids in discrete element modelling. International Journal for Numerical and Analytical Methods in Geomechanics, 19(9):653–659, 1995.

[14] Reza M Baram and Pedro G Lind. Deposition of general ellipsoidal particles. Physical Review E, 85(4):041301, 2012.

[15] Jeffrey D Amack and M Lisa Manning. Knowing the boundaries: extending the differential adhesion hypothesis in embryonic cell sorting. Science, 338(6104):212–215, 2012.

[16] QJ Zheng, ZY Zhou, and AB Yu. Contact forces between viscoelastic ellipsoidal particles. Powder technology, 248:25–33, 2013.

[17] Kamyar Kildashti, Kejun Dong, Bijan Samali, Qijun Zheng, and Aibing Yu. Evaluation of contact force models for discrete modelling of ellipsoidal particles. Chemical Engineering Science, 177:1–17, 2018.

[18] Raymond W Ogden. Non-linear elastic deformations. Courier Corporation, 1997.

[19] X Li, A K Upadhyay, A J Bullock, T Dicolandrea, J Xu, R L Binder, M K Robinson, D R Finlay, K J Mills, C C Bascom, C K Kelling, R J Isfort, J W Haycock, S MacNeil, and R H Smallwood. Skin stem cell hypotheses and long term clone survival–explored using agent-based modelling. Sci. Rep., 3:1904, 2013.

[20] E. Clayton, D. P. Doupe, A. M. Klein, D. J. Winton, B. D. Simons, and P. H. Jones. A single type of progenitor cell maintains normal epidermis. Nature, 446(7132):185–9, 2007.

[21] P. Rompolas, K. R. Mesa, K. Kawaguchi, S. Park, D. Gonzalez, S. Brown, J. Boucher, A. M. Klein, and V. Greco. Spatiotemporal coordination of stem cell commitment during epidermal homeostasis. Science, 352(6292):1471–4, 2016. Rompolas, Panteleimon Mesa, Kailin R Kawaguchi, Kyogo Park, Sangbum Gonzalez, David Brown, Samara Boucher, Jonathan Klein, Allon M Greco, Valentina eng 5R01AR063663-04/AR/NIAMS NIH HHS/ 1R01AR067755-01A1/AR/NIAMS NIH HHS/ T32 GM007223/GM/NIGMS NIH HHS/ P30 AR053495/AR/NIAMS NIH HHS/ T32GM007223/GM/NIGMS NIH HHS/ R01 AR063663/AR/NIAMS NIH HHS/ R01 AR067755/AR/NIAMS NIH HHS/ Research Support, N.I.H., Extramural Research Support, Non-U.S. Gov’t New York, N.Y. 2016/05/28 06:00 Science. 2016 Jun 17;352(6292):1471–4. doi: 10.1126/science.aaf7012. Epub 2016 May 26.

[22] A. Roshan, K. Murai, J. Fowler, B. D. Simons, V. Nikolaidou-Neokosmidou, and P. H. Jones. Human keratinocytes have two interconvertible modes of proliferation. Nat Cell Biol, 18(2):145–56, 2016.

[23] P. Greulich and B. D. Simons. Dynamic heterogeneity as a strategy of stem cell self-renewal. Proc Natl Acad Sci U S A, 113(27):7509–14, 2016.

[24] G. Mascre, S. Dekoninck, B. Drogat, K. K. Youssef, S. Brohee, P. A. Sotiropoulou, B. D. Simons, and C. Blanpain. Distinct contribution of stem and progenitor cells to epidermal maintenance. Nature, 489(7415):257–62, 2012. Mascre, Guilhem Dekoninck, Sophie Drogat, Benjamin Youssef, Khalil Kass Brohee, Sylvain Sotiropoulou, Panagiota A Simons, Benjamin D Blanpain, Cedric eng 079249/Wellcome Trust/United Kingdom 092096/Wellcome Trust/United Kingdom Research Support, Non-U.S. Gov’t England 2012/09/04 06:00 Nature. 2012 Sep 13;489(7415):257–62. doi: 10.1038/nature11393.

[25] Yekaterina A Miroshnikova, Huy Q Le, David Schneider, Torsten Thalheim, Matthias Rübsam, Nadine Bremicker, Julien Polleux, Nadine Kamprad, Marco Tarantola, Irène Wang, et al. Adhesion forces and cortical tension couple cell proliferation and differentiation to drive epidermal stratification. Nature cell biology, 20(1):69, 2018.

[26] M. P. Adams, D. G. Mallet, and G. J. Pettet. Towards a quantitative theory of epidermal calcium profile formation in unwounded skin. PLoS One, 10(1):e0116751, 2015. Adams, Matthew P Mallet, Daniel G Pettet, Graeme J eng Research Support, Non-U.S. Gov’t 2015/01/28 06:00 PLoS One. 2015 Jan 27;10(1):e0116751. doi: 10.1371/journal.pone.0116751. eCollection 2015.

[27] J. L. Maitre, H. Berthoumieux, S. F. Krens, G. Salbreux, F. Julicher, E. Paluch, and C. P. Heisenberg. Adhesion functions in cell sorting by mechanically coupling the cortices of adhering cells. Science, 338(6104):253–6, 2012. Maitre, Jean-Leon Berthoumieux, Helene Krens, Simon Frederik Gabriel Salbreux, Guillaume Julicher, Frank Paluch, Ewa Heisenberg, Carl-Philipp eng Research Support, Non-U.S. Gov’t New York, N.Y. 2012/08/28 06:00 Science. 2012 Oct 12;338(6104):253–6. doi: 10.1126/science.1225399. Epub 2012 Aug 23.

[28] Rete pegs. https://en.wikipedia.org/wiki/Rete_pegs. Accessed: 2019-08-28.

[29] Gary L Grove and Albert M Kligman. Age-associated changes in human epidermal cell renewal. Journal of Gerontology, 38(2):137–142, 1983.

[30] M. Geerligs, L. van Breemen, G. Peters, P. Ackermans, F. Baaijens, and C. Oomens. In vitro indentation to determine the mechanical properties of epidermis. J Biomech, 44(6):1176–81, 2011. Geerligs, Marion van Breemen, Lambert Peters, Gerrit Ackermans, Paul Baaijens, Frank Oomens, Cees eng 2011/02/08 06:00 J Biomech. 2011 Apr 7;44(6):1176–81. doi: 10.1016/j.jbiomech.2011.01.015.

